# Modeling Mechanical Activation of Macrophages During Pulmonary Fibrogenesis for Targeted Anti-Fibrosis Therapy

**DOI:** 10.1101/2023.07.19.549794

**Authors:** Ying Xu, Linxuan Ying, Jennifer K Lang, Boris Hinz, Ruogang Zhao

## Abstract

Pulmonary fibrosis, as seen in idiopathic pulmonary fibrosis (IPF) and COVID-induced pulmonary fibrosis, is an often-fatal lung disease. Increased numbers of immune cells such as macrophages were shown to accumulate in the fibrotic lung, but it is unclear how they contribute to the development of fibrosis. To recapitulate the macrophage mechanical activation in the fibrotic lung tissue microenvironment, we developed a fibrotic microtissue model with cocultured human macrophages and fibroblasts. We show that profibrotic macrophages seeded on topographically controlled stromal tissue constructs become mechanically activated. The resulting co-alignment of macrophages, collagen fibers and fibroblasts promote widespread fibrogenesis in micro-engineered lung tissues. Anti-fibrosis treatment using pirfenidone disrupts the polarization and mechanical activation of profibrotic macrophages, leading to fibrosis inhibition. Pirfenidone inhibits the mechanical activation of macrophages by suppressing integrin αMβ2 (CD11b/CD18) and Rho-associated kinase 2, which is a previously unknown mechanism of action of the drug. Together, these results demonstrate a potential pulmonary fibrogenesis mechanism at the tissue level contributed by mechanically activated macrophages. We propose the coculture, force-sensing microtissue model as a powerful tool to study the complex immune-stromal cell interactions and the mechanism of action of anti-fibrosis drugs.

## Introduction

Pulmonary fibrosis, as seen in idiopathic pulmonary fibrosis (IPF) and COVID-induced pulmonary fibrosis, is a serious and often fatal lung disease [1, 2]. Pulmonary fibrosis is characterized by the significant changes in the architecture, composition and stiffness of the lung tissue that lead to the deterioration of lung functions [3]. Pulmonary fibrosis initiates as a result of alveolar epithelial injury, followed by the recruitment and activation of immune cells and ensuing aberrant wound healing [4]. Increased accumulation of macrophages, including unique pro-fibrotic sub-populations, have been reported in the fibrotic lungs [5]. However, the contribution of different macrophage activation states to the disease progression and severity is still unclear [6]. For instance, macrophages can either contribute to normal repair or attain pro-fibrotic roles, depending on timing, context, and their activation status [7, 8]. The absence of macrophages during inflammation has been shown to result in impaired wound healing [9, 10] while macrophage persistence beyond acute repair phases can result in organ fibrosis [11-13]. Recent studies suggest that the latter effect is at least in part due to the contribution of pro-fibrotic macrophages to the activation of fibroblasts into to myofibroblasts in a process that depends on physical contact [14, 15]. Myofibroblasts are main effector cells in fibrogenesis by producing excessive amounts of collagen and generating high contractile forces by incorporating α-smooth muscle actin (α-SMA) into stress fibers [16]. However, it is still unclear how macrophage-myofibroblast communication at the tissue level contributes to lung tissue fibrogenesis.

In the fibrotic lung tissue, the excessively accumulated fibrillar collagen is often aligned in dense bundles, as a result of the active extracellular matrix (ECM) remodeling by contractile myofibroblasts [17]. This pathologically stiff tissue microenvironment also provides topographic cues to the embedded cells, causing changes in the cellular functions such as their morphology and alignment [18, 19]. For instance, bone marrow-derived macrophages have been shown to align *in vitro* with engineered topographic cues such as microgrooves produced in 2D elastic substrates; alignment promotes profibrotic polarization into pro-fibrotic macrophages characterized by expression of CD206 [20]. For simplicity, *in vitro* generated CD206-positive macrophages are often called M2 phenotypes, recognizing that macrophages exist in a more complex polarization (activation) spectrum *in vivo* [5, 21-26]. Whether and how mechanical and topographical properties of fibrotic tissue microenvironment affect macrophage activation into different polarization states and whether macrophage mechanosensing plays a role in the progression of fibrosis is largely unknown.

Their involvement in fibrogenesis makes immune cells attractive targets for anti-fibrosis therapies [27]. Pirfenidone (PFD) is an FDA-approved anti-fibrotic drug known to act on multiple fibrogenic pathways [28]. PFD reduces fibroblast proliferation and inhibits transforming growth factor-beta (TGF-β)-pathway [29]. Recent studies have shown that PFD can also modulate the polarization and fibrogenic activities of macrophages [30, 31], but since these studies were performed in animal models, it is unknown whether such findings can be applied to humans. In fact, lack of human-relevant, disease-mimetic research models has been one of the major barriers in the development of anti-fibrosis drugs.

Through histological and single cell sequencing analyses of human fibrotic lung samples, we here show that profibrotic macrophages are mechanically activated in fibrotic lung microenvironment. We developed fibrotic microtissues with cocultured human macrophage and lung fibroblasts to model mechanical activation of macrophages and subsequent fibrogenesis. Fibroblast-populated, membranous lung microtissues were first fabricated on a group of micropillars to guide the alignment of collagen fibers and fibroblasts. Profibrotic macrophages seeded onto this tissue became mechanically activated, and their extensive co-alignment with collagen fibers and fibroblasts promoted widespread fibrogenesis in the lung microtissue. PFD treatments disrupted the polarization and mechanical activation of M2 macrophages, leading to fibrosis inhibition. We showed that PFD inhibited macrophage mechanical activation by suppressing integrin αMβ2 (CD11b/CD18) and Rho-associated kinase 2 (ROCK2), which is a previously unknown mechanism of action of PFD. Together, these results reveal a new mechanism how mechanically activated macrophages contribute to pulmonary fibrogenesis at the tissue level. The heterocellular force-sensing microtissue model is shown as a powerful tool to study the complex immune-stromal cell interactions and the mechanism of action of anti-fibrosis drugs.

## Results

### Mechanical activation of lung macrophages in fibroblast-remodeled fibrotic tissue microenvironment

To spatially resolve the interaction of collagen, macrophages, and fibroblasts in human fibrotic lungs, we processed lung tissues from donors suffering from IPF for histological analysis. H&E staining of highly fibrotic regions revealed a significantly remodeled tissue architecture, characterized by large amounts of elongated cells embedded between dense and aligned ECM fibers (Fig. 1A, Supplementary Fig. 1). The embedded cells consisted of a large number of α-SMA-positive myofibroblasts and CD206-positive macrophages that were in proximity. Myofibroblasts and macrophages both adopted elongated morphologies and aligned with each other, suggesting they sensed and responded to the remodeled tissue microenvironment (Fig. 1B, Supplementary Fig. 1). Re-analysis of the dataset from a single-cell RNA-sequencing (scRNA-seq) study of human donor samples revealed a unique sub-population of profibrotic alveolar macrophages that were exclusively present in patients with lung fibrosis (Fig. 1C) [32]. Our analysis showed elevated expression levels of genes associated with mechanotransduction pathways in these alveolar macrophages, including *RHOA, ITGB2*, and *ITGAM*, compared to macrophages obtained from healthy donors (Fig. 1D-H). These analyses suggest that lung macrophages undergo mechanical activation in the highly remodeled fibrotic tissue microenvironment of human IPF lungs.

**Fig. 1.**
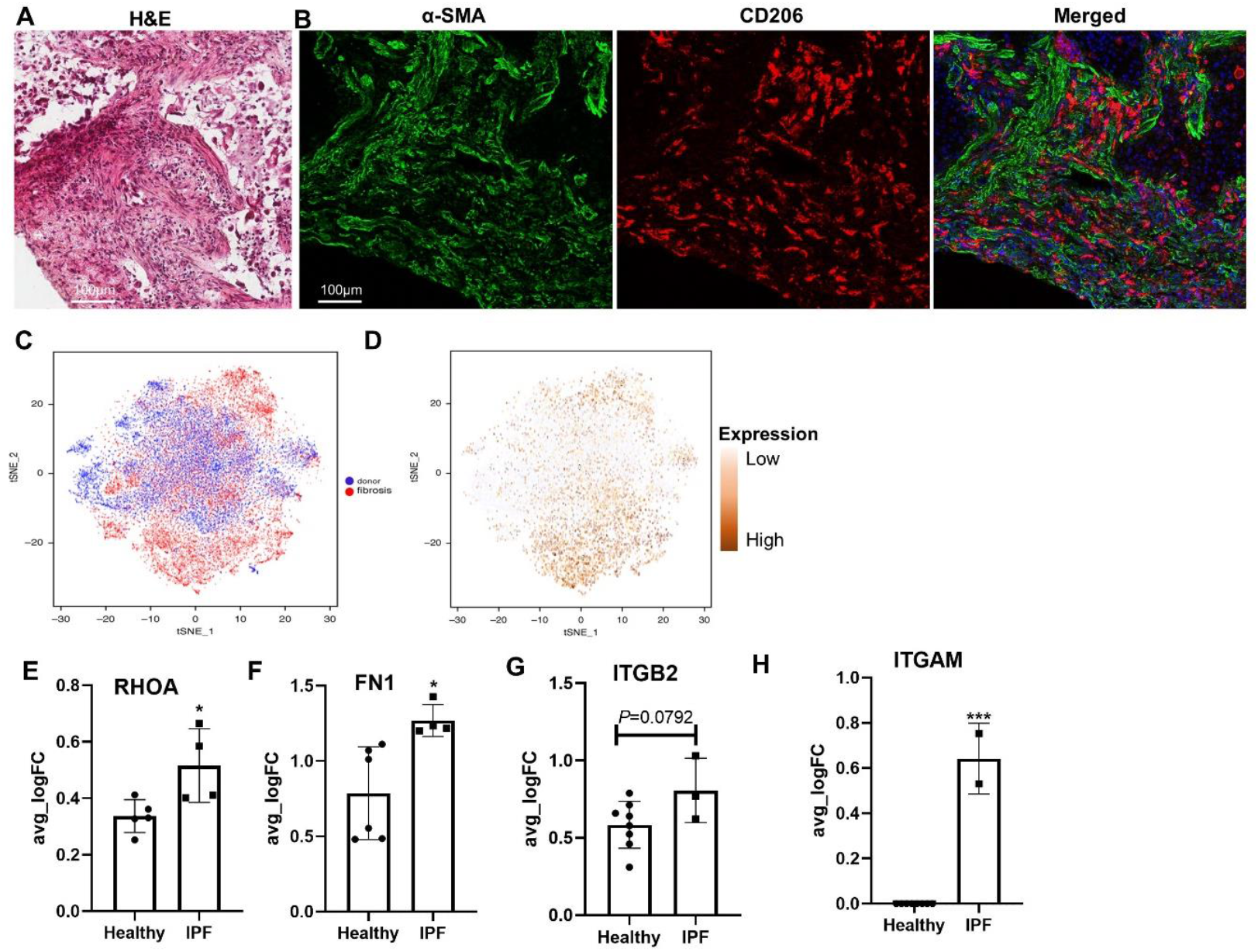
Mechanical activation of lung macrophages in a highly remodeled fibrotic tissue microenvironment. (A) H&E staining of a highly remodeled region in a fibrotic lung tissue. (B) Confocal fluorescence images of α-SMA and CD206 of a matching region in an adjacent tissue section. Both α-SMA^+^ myofibroblasts and CD206^+^ macrophages adopted elongated morphology and aligned with each other. Scale bars, 100 μm. (C) t-SNE plot showing the distinct populations of alveolar macrophages in IPF patients and healthy donors. (D) t-SNE plot showing the preferential expression of ITGAM gene in alveolar macrophages in the IPF patients illustrated in panel C. Data extracted from healthy subjects (n=8) and IPF patients (n=8) based on NCBI GEO Dataset: GSE122960. Plots of averaged gene expression for RHOA (E), FN1 (F), ITGB2 (G) and ITGAM (H) in alveolar macrophages from healthy and IPF donor lungs, based on Dataset: GSE122960.

### Macrophages are mechanically responsive to topography-controlled tissue remodeling

To understand how macrophages respond to remodeled tissue architecture such as those seen in fibrotic lung tissue, we created engineered microtissues with controlled morphology and cell and ECM alignment. Normal human lung fibroblasts (NHLF) were mixed with collagen type I and seeded into microfabricated microwells containing a group of micropillars arranged in topological patterns. Pattern designs included a spiral pattern (Fig. 2A), diamond pattern (Fig. 2B) and square pattern (Fig. 2C) to mimic various geometries found in the lung tissue such as highly curved alveolar wall. Fibroblasts generated contractile forces and compacted the collagen ECM against the micropillars. As a result of fibroblast-mediated ECM remodeling, morphologically controlled microtissues formed following the micropillar array pattern templates. NHLF and collagen fibers in the microtissues were strongly co-aligned along the lines of tension defined by the micropillar boundary conditions (Supplementary Fig. 2). Thus, controlled cell and ECM fiber alignment provide an opportunity to examine macrophage response to the tissue topography observed in fibrosis. Next, monocytes derived from human peripheral blood were pre-polarized with interleukin (IL)-4 and IL-13 before being seeded onto NHLF-remodeled microtissues. Such M2 macrophages adhered to the microtissue surface within 48 h and, remarkably, adopted an elongated spindle morphology aligning with collagen fibers and NHLF (Fig. 2 D-F, J, K, and Supplementary Fig. 2). Confocal imaging revealed that macrophages at least partially penetrated the microtissue and interacted with the surrounding collagen fibers and NHLF in 3D, as demonstrated by the overlapping collagen fibers and macrophages (Fig. 2 G-I). These results suggest that M2 macrophages are capable of sensing topographic cues in fibrosis-like tissue microenvironments and actively adapt to these cues.

**Fig. 2.**
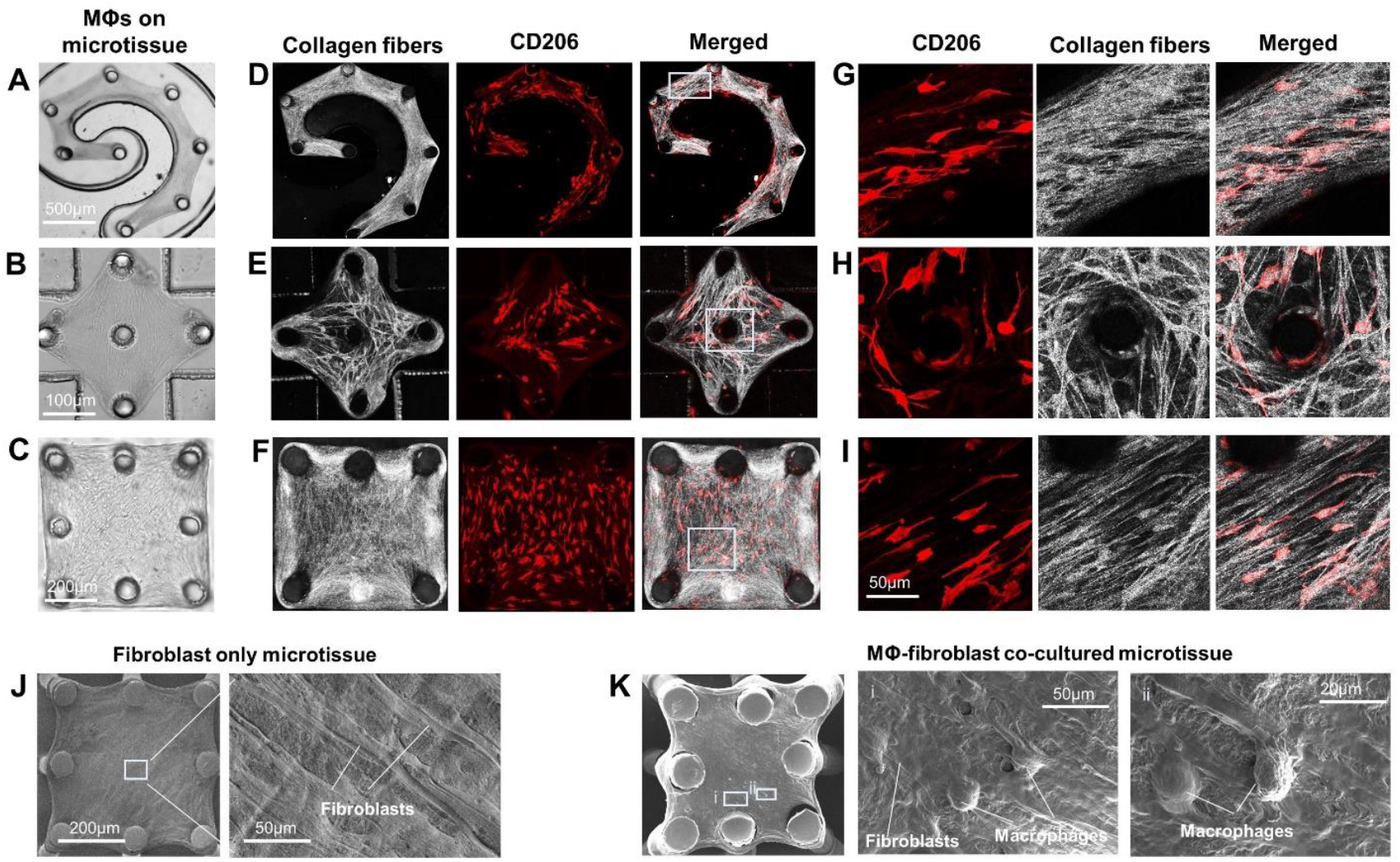
Macrophage mechanical activation in response to topographically-controlled tissue remodeling. Phase contrast images of fibroblast and macrophage cocultured lung microtissues formed on a group of flexible micropillars arranged in a spiral pattern (A), a diamond pattern (B) and a square pattern (C). Confocal reflectance images of collagen fibers (gray) and fluorescence images of CD206 (red) of spiral patterned (D), diamond patterned (E) and square patterned (F) microtissues. (G, H, I) Enlarged views of boxed regions in (D-F) showing the co-alignment between the CD206-positive macrophages and the collagen fibers. (J) SEM images of a fibroblast only microtissue showing the fibroblast alignment. (K) SEM images of a cocultured microtissue showing the co-alignment between fibroblasts and macrophages.

### Extensive co-alignment between macrophages and fibroblasts promotes a widespread fibrosis response in engineered microtissues

Next, we sought to understand how profibrotic macrophages contribute to the formation of fibrosis in the macrophage-NHLF coculture microtissues. Profibrotic M2 macrophages were introduced to the surface of NHLF-populated microtissues (Fig. 3A). After 3 d of coculture, robust expression of α-SMA was observed in the microtissues, indicating NHLF activation into myofibroblasts (Fig. 3B). The expression levels of α-SMA in M2 macrophage cocultured microtissues were significantly higher (4.7 fold) than those in NHLF-only microtissue, but moderately lower than TGF-β1 treated positive controls (Fig. 3B, C). Since differentiated myofibroblasts have been shown to generate high contractile forces, we measured the contractile force generation in monocultured and cocultured microtissues using in-situ micropillar force sensors. M2 macrophage and NHLF cocultured microtissue was shown to generate 5.7 times higher contractile force as compared to NHLF monocultured microtissues (Fig. 3D). Concomitantly, levels of active TGF-β measured in the culture supernatant were highest in the coculture conditions (Fig. 3E), suggesting a potential role of paracrine TGF-β signaling in macrophage-mediated tissue fibrosis. Since early studies have shown that activation of fibroblasts by macrophages mainly occurs in close proximity [15], we examined co-registration of both cells in the cocultured microtissue. Corresponding to the patterns observed in fibrotic lung tissues (Fig. 1A, Supplementary Fig. 1), elongated CD206-positive M2 macrophages abundantly aligned with α-SMA-positive myofibroblasts throughout the whole microtissue (Fig. 3B, merged). At high microscopy resolution, single macrophages were in proximity and/or direct contact with myofibroblasts (Fig. 3F).

**Fig. 3.**
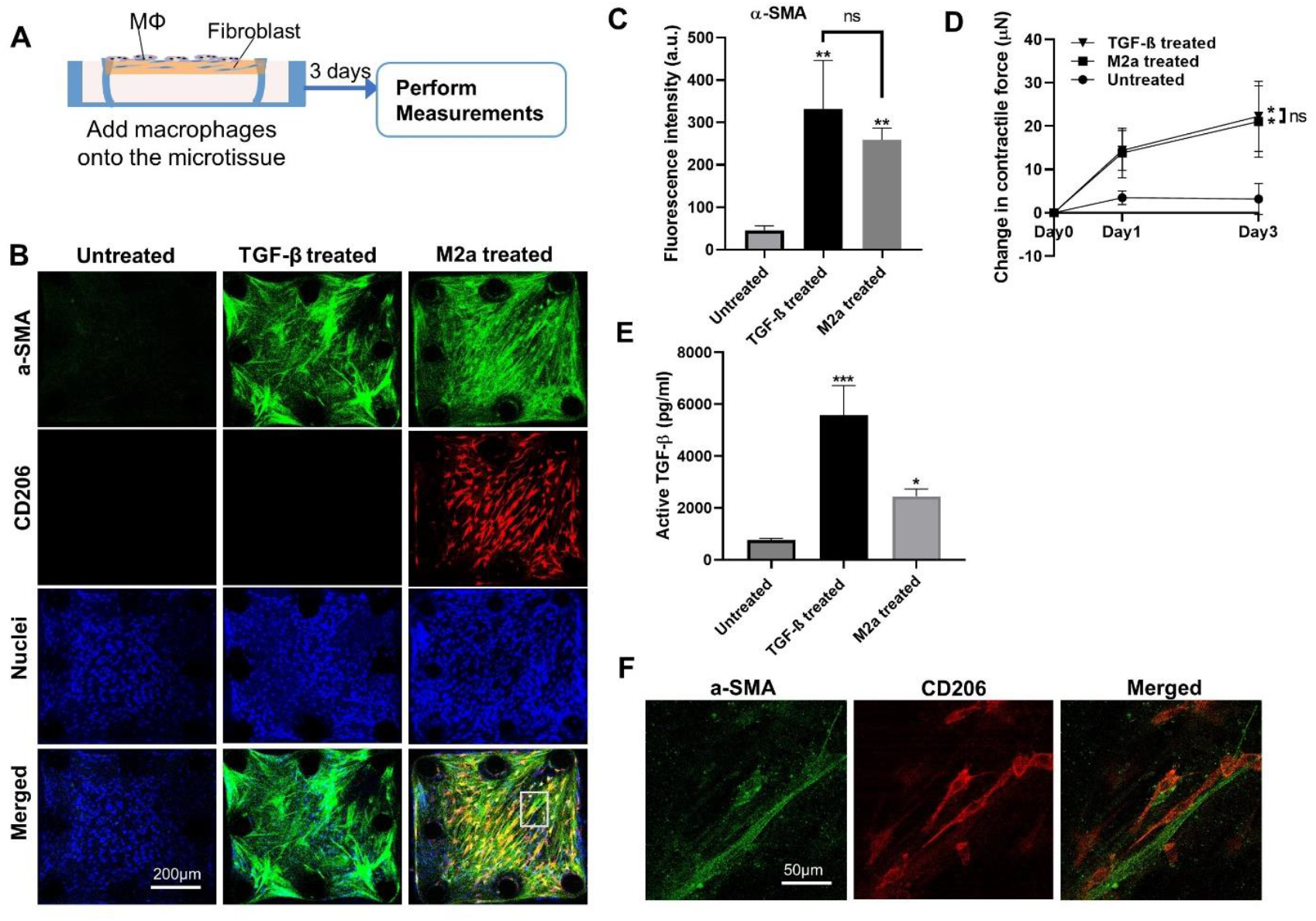
Extensive co-alignment between macrophages and fibroblasts promoted a widespread fibrosis response in the microtissue. (A) Schematic showing the fibrosis induction in NHLF-populated microtissues by pro-fibrotic macrophages. (B) Representative fluorescence images of NHLF only (untreated), TGF-β1 treated NHLF only, and M2 macrophage and NHLF cocultured microtissues (left to right). Microtissues were stained for nuclei (blue), α-SMA (green) and CD206 (red). Scale bar is 200 μm. (C) Fluorescence intensity measurement of α-SMA for different microtissue culture conditions. (D) Contractile force measurement for different microtissue culture conditions. (E) Supernatant active TGF-β level for different microtissue culture conditions. (F) Enlarged view of the boxed region in panel B showing the co-alignment between single myofibroblasts and M2 macrophages.

### Anti-fibrosis treatment with PFD acts on macrophage-induced fibrosis

PFD and Nintedanib are the only FDA-approved drugs for the treatment of lung fibrosis; both with unclear modes of action and cellular targets in the fibrotic tissue [33]. Our macrophage-NHLF cocultured microtissue (MaFiCo) is an excellent fibrosis model to interrogate the action of anti-fibrotic drugs. As a paradigm drug, we applied PFD to our microtissue model using (1) preventative treatment, (2) therapeutic treatment, and (3) pre-treatment. The purpose of these different regimens was to work either without pre-existing fibrosis condition (preventative treatment), or with pre-existing fibrosis condition (therapeutic treatment). Pre-treatment was performed only on macrophages to decouple PFD’s effect on the two cell types.

In the preventative treatment, PFD was introduced together with the macrophages to the NHLF-populated microtissues and maintained in the coculture for 3 d (Fig. 4A). Preventative PFD treatment significantly reduced the number of adhered macrophages by 86% (Fig. 4B,C), α-SMA expression in NHLFs (95%) (Fig. 4B, D), microtissue contractile forces (144%) (Fig. 4E), and active TGF-β levels (62%) (Fig. 4F) compared to untreated controls in a dose-dependent manner. At the highest concentration of 1000 μg/ml, PFD almost entirely abolished macrophage adhesion to the microtissue surface and expression of α-SMA in NHLF (Fig. 4B). The few remaining macrophages in this condition exhibited super elongated shapes, which were drastically different from the spindle shapes attained in untreated controls (Fig. 4B). Since the microtissues did not have pre-existing fibrotic features in the preventative treatment, these results suggest that disruption of macrophage adhesion to the microtissues inhibited the development of fibrosis.

**Fig. 4.**
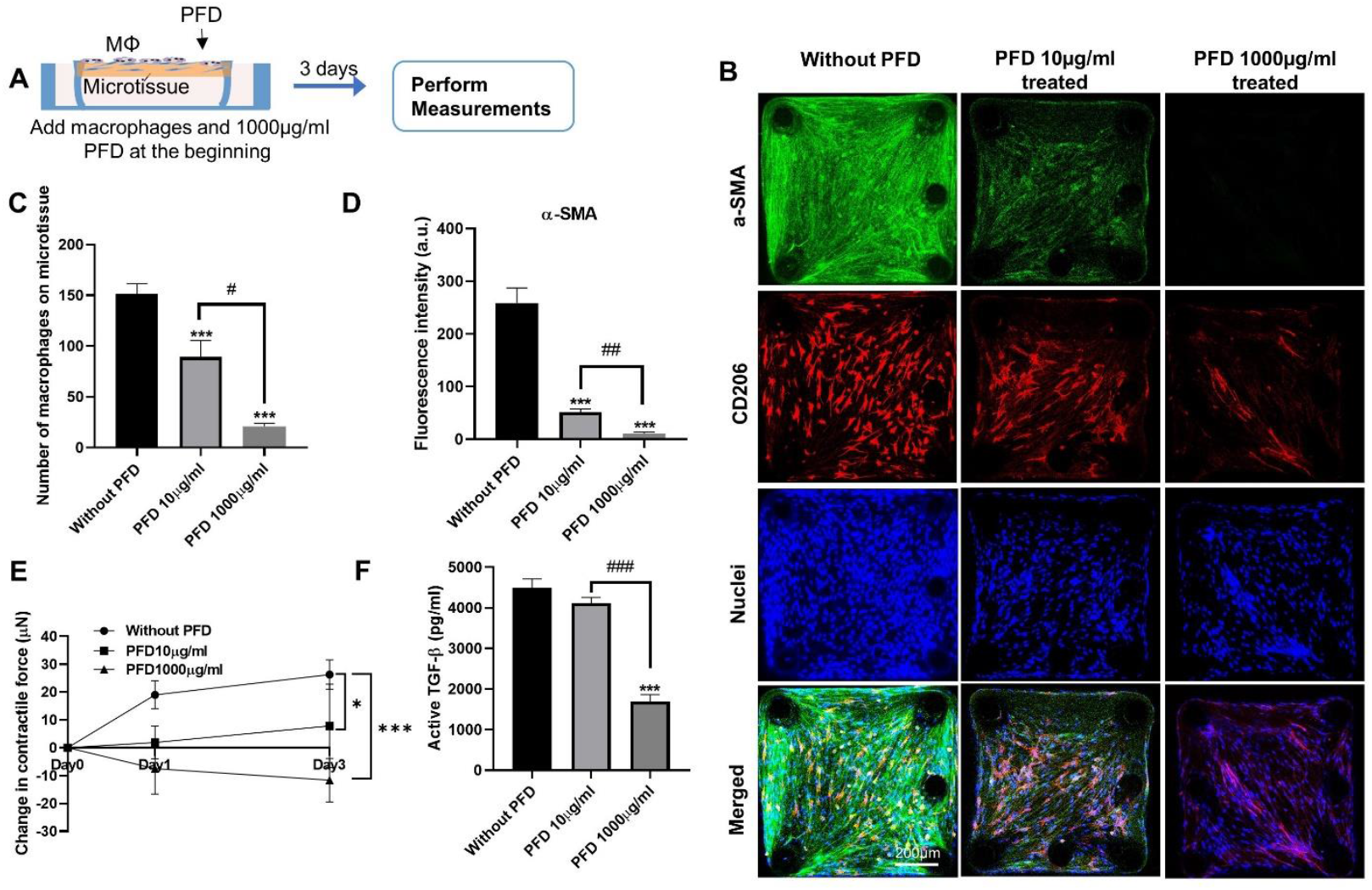
The effect of preventative anti-fibrosis treatment on cocultured microtissue. (A) Schematic showing the preventative anti-fibrosis treatment and results evaluation. Pirfenidone (PFD) was administrated at the beginning of the coculture. (B) Representative fluorescence images of cocultured microtissues treated with 0, 10 or 1000 μg/ml PFD (left to right). Scale bar is 200 μm. (C) The number of macrophages adhered on the microtissue under different PFD systems. (D) Fluorescence intensity measurement of α-SMA under different PFD treatment conditions. (E) Contractile force measurement under different PFD treatment conditions. (F) Supernatant active TGF-β level under different PFD treatment conditions.

In the therapeutic treatment regimen, macrophages and NHLF were cocultured for 3 d to form fibrotic microtissues before PFD was introduced for another 3 d (Fig. 5A). PFD treatment only moderately reduced the numbers of attached macrophages (40%) (Fig. 5 B,C), α-SMA expression in NHLFs (47.5%) (Fig. 5B, D) and levels of active TGF-β in the supernatant (Fig. 5F), as compared to the untreated control microtissues. The change in the microtissue contractile forces under PFD treatment is not significant (Fig. 5E). Furthermore, PFD treatment caused a significant change in the morphology of adhered macrophages from spindle to thread shapes (Fig. 5B), as seen in the preventative treatment. However, compared to the preventative treatment (Fig. 4C), more macrophages remained adherent in the therapeutic treatment (Fig. 5C).

**Fig. 5.**
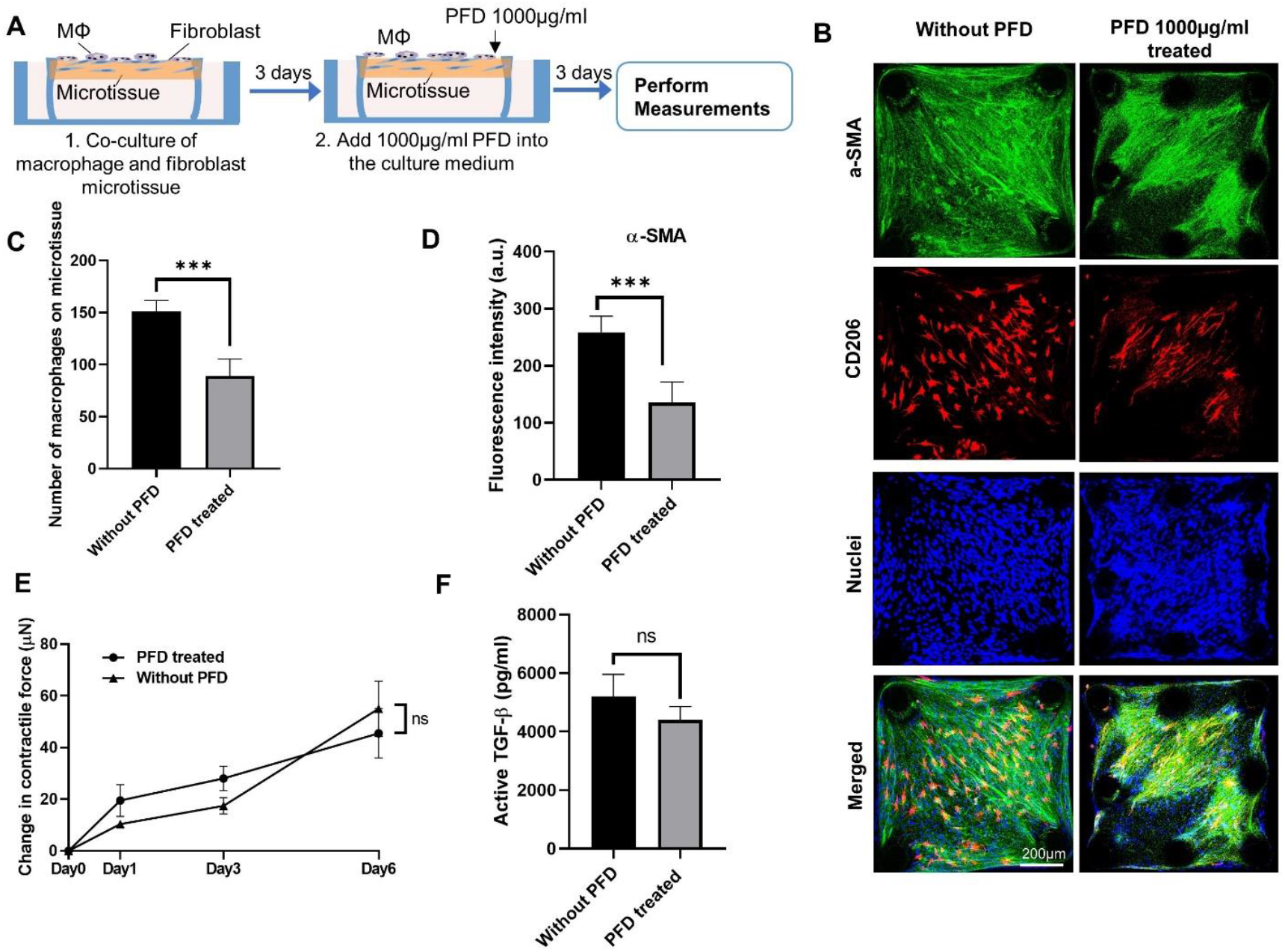
The effect of therapeutic anti-fibrosis treatment on cocultured microtissue. (A) Schematic showing the therapeutic anti-fibrosis treatment and results evaluation. Pirfenidone (PFD) was administrated after 3 d of coculture. (B) Representative fluorescence images of cocultured microtissues treated with 1000 μg/ml PFD. Scale bar is 200 μm. (C) The number of macrophages adhered on the microtissue under different PFD systems. (D) Fluorescence intensity measurement of α-SMA under different PFD treatment conditions. (E) Contractile force measurement under different PFD treatment conditions. (F) Supernatant active TGF-β level under different PFD treatment conditions.

Since PFD treatment affected both macrophages and NHLF, we next pre-treated macrophages with PFD to decouple its effect on these two cell types. Human peripheral blood-derived monocytes were first treated with M-CSF, followed by addition of PFD for 24 h before inducing M2 polarization with IL-4/IL-13 for another 24 h (Fig. 6A). Adding PFD during induced M2 polarization inhibited the expression of CD206 and reduced active TGF-β levels measured in the macrophage monoculture supernatants in a dose-dependent manner (Supplementary Fig. 5). Such pre-treated macrophages were then seeded onto NHLF-populated microtissues and maintained without further addition of PFD. After 3 d coculture, there were significant reductions in the number of adhered macrophages by 72% (Fig. 6 B,C), α-SMA expression in NHLFs (93%) (Fig. 6B, D), microtissue contractile forces (100%) (Fig. 6E) and levels of active TGF-β by 50% (Fig. 6F) compared to untreated controls in a dose-dependent manner. Together, these results showed that PFD inhibits both induced M2 polarization of the macrophages and their capability to adhere to NHLF-populated microtissues. Since both functions are related to macrophage-mediated fibrogenesis, PFD may, in fact, act on macrophages to affect their contribution to the fibrosis process.

**Fig. 6.**
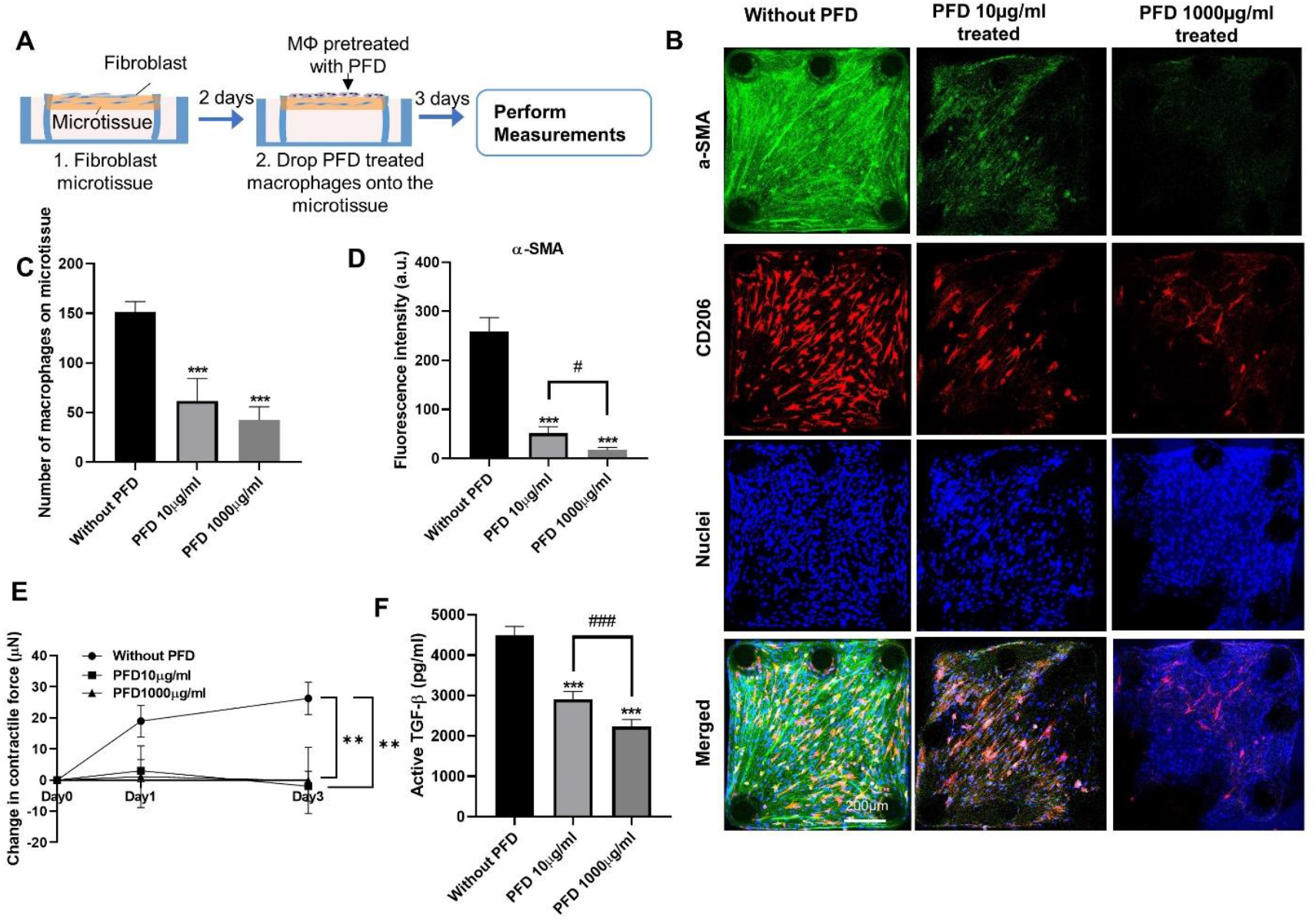
The effect of anti-fibrosis pre-treatment on cocultured microtissue. (A) Schematic showing the anti-fibrosis pre-treatment and results evaluation. Macrophages were pre-treated with PFD for 24 h and then added to NHLF microtissues to form the coculture. (B) Representative fluorescence images of cocultured microtissues treated with 0, 10 or 1000 μg/ml PFD (left to right). Scale bar is 200 μm. (C) The number of macrophages adhered on the microtissue under different PFD systems. (D) Fluorescence intensity measurement of α-SMA under different PFD treatment conditions. (E) Contractile force measurement under different PFD treatment conditions. (F) Supernatant active TGF-β level under different PFD treatment conditions.

### PFD suppresses mechanical activation of macrophages through inhibition of ROCK2 and integrin αMβ2

Treatment with PFD reduced the ability of macrophages to attach, align, and spread on microtissues, which are all cell functions dependent on cell adhesion. In turn, M2 polarization has been shown to require strong adhesion and intracellular contractile forces ([34]). Concomitantly, we identified elevated gene expression of RhoA (contraction) and macrophage integrins αM and β2 (adhesion) in published scRNAseq datasets of lung macrophages of IPF patients (Fig. 1) [35]. Thus, we focused on RhoA-related pathways, αM integrin, and β2 integrin to identify whether PFD suppresses fibrogenesis by acting on macrophage mechanosensing and activation. Rho-associated kinases ROCK1 and ROCK2 are major downstream effectors of the RhoA and ROCK2 has been used as a therapeutic target in clinical trials of anti-fibrosis drug KD025 [36]. Immunostaining of ROCK2 in the coculture microtissues show that ROCK2 predominantly expressed in the M2 macrophages. Treatment with PFD caused 65% and 74% reduction in the expression of the ROCK2 in both preventative and therapeutic treatment regimens (Fig. 7A-C). Next, we treated the coculture microtissue with KD025, a ROCK2 specific inhibitor, to determine whether the suppression of macrophage mechanical activation is ROCK2 specific. KD025 was shown to cause 39% and 21% reductions in the expression of the ROCK2 in preventative and therapeutic treatments, respectively (Fig. 7A-C), together with significant reduction in the adhered macrophage numbers (75% in preventative and 67% in therapeutic) (Fig. 7D,E), significant change in the macrophage morphology from spindle to thread shape and significant reduction in the active TGF-β (62% in preventative and 15% in therapeutic) (Fig. 7F, G). These effects are very similar to those of the PFD treatment. To determine whether the above effects are specific to the macrophages, both PFD and KD025 treatment were performed on monoculture M2 macrophages on silicone substrate (stiffness 4MPa). ROCK2 expression was significantly reduced by PFD treatment (71%) and KD025 treatment (70%), together with 41% and 67% reductions in the number of attached macrophages and 40% and 35% reductions in the active TGF-β (Supplementary Fig. 6). These effects are similar to those in the coculture, suggesting the inhibition of ROCK2 pathway is specific to the M2 macrophages.

**Fig. 7.**
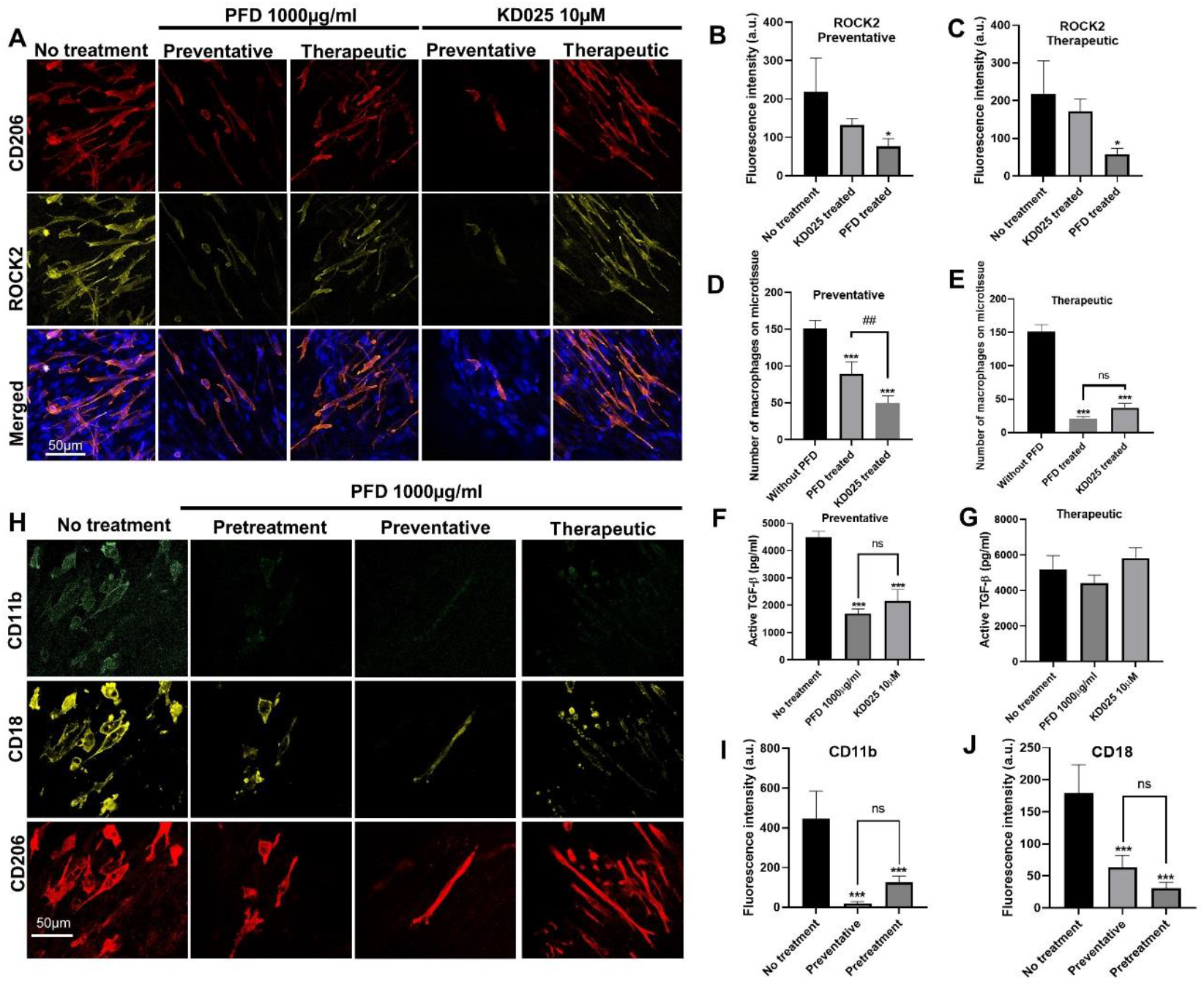
PFD suppresses macrophage mechanical activation through inhibition of ROCK2 and integrin αMβ2. (A) Representative immunofluorescence images of macrophage and fibroblast cocultured microtissues under PFD or ROCK2 inhibitor KD025 treatments. Microtissues were stained for CD206 (red), ROCK2 (yellow) and Nuclei (blue) from top to bottom. Fluorescence intensity measurement of ROCK2 in cocultured microtissues under preventative treatment (B) and therapeutic treatment (C). Number of adherent macrophages on cocultured microtissues under preventative treatment (D) and therapeutic treatment (E). Active TGF-β in cell culture supernatant under preventative treatment (F) and therapeutic treatment (G). (H) Representative immunofluorescence images of macrophage and fibroblast cocultured microtissues under different PFD treatment conditions. Microtissues were stained for CD11b (green), CD18 (yellow) and CD206 (red) from top to bottom. Fluorescence intensity measurement of CD11b (I) and CD18 (J) under treatments with or without PFD.

We next examined the expression of αM and β2 integrins in coculture microtissues. Immunostaining showed that these integrins exclusively expressed in the M2 macrophages. Treatment with PFD caused 64% and 83% reductions in the expression of the αM and β2 integrin subunits in both preventative and pretreatment regimens (Fig. 7H-I). To determine whether this effect is specific to the macrophages, PFD treatment was performed on M2 macrophages in monocultures. Expression of αM and β2 integrins was also significantly reduced by PFD treatment in the monoculture (65% and 96%) (Supplementary Fig. 7). Since RhoA/ROCK signaling affects the expression of integrins, we also treated the M2 macrophage monoculture using KD025. ROCK2 inhibition by KD025 significantly reduced expression of the αM (81%) and β2 (60%) integrins (Supplementary Fig. 7), similar to the effect of PFD treatment. Together, these results show that PFD has inhibitory effects on the macrophage mechanotransduction pathways including the ROCK2 and αM and β2 integrins, and these effects are similar to those of the selective ROCK2 inhibitor KD025.

## Discussion

Pulmonary fibrosis is a deadly disease, but its disease mechanism is still unclear. The involvement of the M2 profibrotic macrophages in the pulmonary fibrogenesis has been broadly reported based on the bulk tissue analysis of the cytokine release and the single cell RNA sequencing analysis [32, 37]; however, these studies do not provide a clear picture of the interaction between macrophages and fibroblasts and remodeled ECMs. As a result, the fibrosis formation process at the tissue and cellular level is still unclear. This has led to difficulties identifying cellular and molecular targets for anti-fibrosis therapies. In the current study, a macrophage and fibroblast coculture fibrosis tissue model was developed to allow the investigation of immune-stroma interactions in a fibrosis microenvironment. Using tissue microfabrication method, we created micropillar-based lung microtissues with controlled topographies to mimic the alignment of fibroblasts and collagen fibers in the fibrotic lung tissue. We showed that profibrotic macrophages were able to sense and adapt to the topographic cues and become elongated and align along the fibroblasts and collagen fibers. At the single cell level, such co-alignment allowed ample contact between the macrophage and the fibroblast. Since it has been shown that proximity is crucial for the crosstalk between macrophages and fibroblasts [ref], the increased level of physical contacts between these two cell types will likely enhance the cellular signal transduction and promote macrophage-mediated myofibroblast differentiation. At the tissue level, the report on the distribution of profibrotic macrophages in lung fibrosis and their relative position to the fibroblasts is limited. Through histological analysis, we showed that CD206^+^ profibrotic macrophages were widely distributed in the fibrotic lung tissue and embedded in the highly deformed ECM along with α-SMA-positive myofibroblasts. The coculture model in the current study recapitulated this effect and demonstrated that extensive co-alignment between the macrophages and fibroblasts enabled the widespread fibrosis response in the tissue, as seen in lung fibrosis.

Lung fibrosis is associated with a mechanically active tissue microenvironment. Differentiated myofibroblasts generate high contractile forces that can cause cell and ECM fiber alignment and disrupt the native alveolar architecture [3, 38, 39]. Macrophages are known to be mechanically sensitive and M2 macrophages have been shown to align on the microgrooves created on 2D elastic substrate [20]. Increased alignment promoted the M2 differentiation of the macrophages. Aligned patterns with feature sizes in the micron level were shown to generate M2 macrophages through forced alignment, whereas irregular topographies with similar dimensions produce circularly spreading macrophages with M1 characteristics [40-46]. In addition to surface topographies, human and mouse lineage and primary macrophages were shown to respond to the elastic modulus (‘stiffness’) of their substrates [47]. In general, macrophages attaching to soft materials exhibit reduced activation into profibrotic M2 macrophages compared to those grown on stiffer substrates [48-52]. Furthermore, macrophages are able to sense and follow changes in the deformation of collagen networks caused by contracting fibroblasts over millimeter distances [53].

Here, we show that fibroblast-mediated cell and ECM fiber alignment provided an instructive microenvironment for the macrophage mechanical activation, which may imply a mechanism of macrophage recruitment and activation in fibrotic diseases. We further showed that mechanical activation of macrophages led to elevated level of TGF-β secretion and myofibroblast differentiation, suggesting that mechanical activation is an important step in the activation of the pro-fibrotic functions of macrophages. Since fibroblast activation and elevated tissue mechanical properties have been shown to form a feedback loop in the fibrogenesis [54], the mechanical activation of macrophages may participate in this feedback loop and contribute to the sustained progression of fibrosis. Given this finding, the mechanical activation of macrophages has the potential to be used as a target for anti-fibrosis therapies.

Current therapies for fibrotic diseases are very limited. PFD and Nintedanib are the only two drugs approved by FDA to treat IPF, but they can only slow down the disease progression and cannot stop or reserve the disease [33]. The mechanisms of action of PFD and nintedanib are not fully understood. PFD is generally believed to act on fibroblasts through inhibiting TGF-β pathway, but recent studies have shown that PFD can also modulate the macrophage polarization and profibrotic activities [30, 31]. In the current study, we showed that PFD strongly affected the M2 macrophage adhesion on the fibroblast-populated stromal tissue through inhibiting the ROCK2 and integrin αMβ2. Since macrophage adhesion is critical to their recruitment to the injury site, this finding suggests a previously unknown mechanism of action of PFD on macrophage-mediated fibrosis. Further histological and single cell studies on PFD treated donor lung samples are needed to determine the clinical relevancy of this finding. If confirmed, this may open a new avenue for the development of anti-fibrosis drugs that target immune cells. The above findings have been enabled by the cocultured, force-sensing microtissue model developed in this study. Compared to the previous cocultured models of immune cells and stromal cells [15, 55-58], the current model has unique features to allow the manipulation of the mechanical microenvironment and the measurement of the change in the tissue mechanical properties. As a result, it provides novel perspectives to the immune-stroma interaction and can unveil previously unknown disease and therapeutic mechanisms. Together, this novel system has the potential to become a powerful tool to study the complex immune-stromal cell interactions and the mechanism of action of anti-fibrosis drugs.

## Materials and Methods

### Peripheral blood monocyte isolation and pro-fibrotic macrophage differentiation

Whole blood was collected at the Clinical and Translational Research Center in Buffalo according to approved protocols with the University at Buffalo Research Subjects Institutional Review Board, and with informed consent obtained from all donors. The PBMCs were isolated through density gradient centrifugation, where 18 mL of blood was mixed with Hanks’ Balanced Salt Solution (HBSS, 14175095, GibcoTM) before being poured onto Ficoll-Paque Plus (GE17-1440-03, GE Healthcare). The layers were separated through centrifugation, and the cells from the interphase were washed twice with HBSS before being resuspended in 5 mL of HBSS for monocyte purification. The classical monocytes were isolated using biotin-conjugated antibodies and streptavidin-conjugated magnetic beads to deplete labeled cells and were then conditioned into resting macrophages (M0) using RPMI 1640 supplemented with 10% fetal bovine serum (FBS, S11150H, R&D Systems) and 1% penicillin/streptomycin (P/S, 15140122, Gibco™) for 7 d. After 4 d, fresh media with 50 ng/ml macrophage colony stimulating factor (M-CSF, PHC9504, GibcoTM) was added, and mature macrophages were polarized by adding 20 ng/ml Interleukin-4 (IL4, A42602, Invitrogen) plus 20ng/ml IL13 (A42526, Invitrogen) to generate M2 macrophages for an additional 24 h. Finally, for monocytes or macrophages cultured on glass coverslip, the culture surface of glass coverslips was coated with rat tail collagen type-I (5153, Advanced Biomatrix) before cell seeding.

### NHLF cell culture and coculture assays

Primary NHLF were purchased from Lonza and subcultured for up to six passages in manufacturer supplied growth medium (FGM-2 BulletKit, CC3132, Lonza). The microtissues populated with NHLF were grown for 2 d before the addition of M2 macrophages at a ratio of 4:1 (M2 : NHLF). Cocultures of macrophages and NHLF were carried out for 3 d in macrophage base medium containing RPMI1640, 10% FBS, and 1% P/S.

### Micropillar device fabrication and microtissue seeding

The fabrication of micropillar arrays involved the utilization of a multi-layer microlithography technique, as previously described [ref]. In summary, the process included the following steps: First, SU-8 masters were created by spin coating multiple layers of SU-8 photoresists, followed by alignment, exposure, and baking. To achieve a cross-sectional difference between the micropillar head and leg sections, a thin layer of SU-8 doped with S1813 was deposited, acting as a barrier to prevent UV light penetration into the leg section. UV exposure was conducted using an OAI mask aligner equipped with a U-360 band pass filter. Subsequently, PDMS (Sylgard 184, DowCorning) stamps were cast over the SU-8 master using a 10:1 mixing ratio. The micropillar devices were then generated through replica molding from the PDMS stamps, employing P35 petri dishes as the mold. For microtissue seeding, a previously established protocol was employed [ref]. The micropillar devices were sterilized and treated with Pluronic F-127 (P2443, Sigma) to prevent non-specific cell adhesion to the PDMS surface. NHLFs were mixed with rat tail collagen type-I (5153, Advanced Biomatrix) at a final concentration of 2 mg/ml. The mixture was introduced into the microwells by centrifugation. Subsequently, the collagen solution was crosslinked and maintained in appropriate growth media within a CO2 incubator. After the microtissue formation, additional NHLF cells were introduced to the surface of the microtissues before loading the macrophages.

For tracing purposes in some conditions, cells were fluorescently labelled. One million human macrophages and NHLF were suspended in 1 mL of serum-free RPMI1640 medium. To label the cells, 5 μL of Vybrant™ DiO Cell-Labeling Solution (V22886, Invitrogen™) was added to the NHLF suspension, while 5 μL of Vybrant™ DiI Cell-Labeling Solution (V22885, Invitrogen™) was added to the macrophage suspension. The cells were then incubated at 37 °C, 95% humidity and 5% CO2 for 10 min before being washed three times with phosphate buffered saline (PBS, 10010023, Gibco™).

### Pharmacological treatment

Anti-fibrotic drugs Pirfenidone (P1871, TCI America) and KD025 (HY-15307, MedChemExpress) were purchased from commercial suppliers. To induce myofibroblast differentiation in a macrophage-free culture system, 5 ng/ml of transforming growth factor beta 1 was added to the macrophage base media and maintained for 3 d. To test the drugs in a therapeutic manner, human monocytes derived M0 macrophages were cultured in macrophage base medium supplemented with M-CSF, IL-4, and IL-13 to promote M2 polarization. After 24 h, macrophages were collected and loaded onto NHLF populated microtissues, which were maintained in macrophage base medium for 3 d to induce fibrosis. On day 3, the culture medium was refreshed with 0, 10, or 1000 μg/ml PFD co-administered into the coculture system, which was maintained for another 3 d.

To test the drugs in a preventative manner, human monocytes derived M0 macrophages were cultured in macrophage base medium supplemented with M-CSF, IL-4, and IL-13 to promote M2 polarization. After 24 h, macrophages were collected and loaded onto NHLF populated microtissues. PFD was co-administered at 0, 10, or 1000 μg/ml into the cocultured system, which was maintained for 3 d in macrophage base medium. To investigate the anti-fibrotic effect of PFD on macrophages, human monocytes derived M0 macrophages were cultured in macrophage base medium supplemented with pirfenidone for 24 h before treatment with cytokines. After another 24 h, M0 macrophages were stimulated with M-CSF and IL-4 plus IL-13 to promote M2 polarization. Then, macrophages were collected and loaded onto normal NHLF populated microtissues, which were maintained in macrophage base medium for 3 d.

### Microtissue contraction force measurement

The force of spontaneous contraction generated by individual microtissues was determined by measuring the deflection of micropillars using cantilever bending theory. The deflection of the micropillars was measured by comparing the deflected position of the centroid of each pillar top to the centroid of its base using phase contrast microscopy. The microtissue contraction force was measured for the same batch of samples over different time points for each pharmacological condition. Image analysis was conducted using ImageJ software, and the absolute values of the calculated force from each micropillar were added together and divided by 2 to obtain the collective force generated by one microtissue.

### Active TGF-β measurements

To quantify the amount of active TGF-β, TMLCs (mink lung epithelial cells) reporter were used. These cells produce luciferase in response to TGF-β under the control of the PAI-1 promoter. To determine the amount of TGF-β in culture supernatants, TMLCs (40,000 cells/cm2) were allowed to adhere for 3 h, after which they were exposed to conditioned media or native (active TGF-β) for 24 h. TMLCs were then lysed and the luminescence was measured using a luciferase assay kit (Promega) and luminometer (Centro LB, Berthold Technologies). All results were adjusted to account for the baseline luciferase production of TMLCs in the absence of TGF-β. TGF-β concentrations were determined using standard curves generated with known concentrations of active TGF-β in serum-free medium (ranging from 0-8,000 pg/ml) (Supplementary Fig. 4).

### Immunofluorescence and image analysis

The human lung cell-populated microtissues were fixed with 3.7% formaldehyde to preserve their cellular structure. Then, cold methanol was used to permeabilize the cells, allowing the antibodies to penetrate and bind to their target proteins. The cells were then blocked with 3% BSA to prevent non-specific binding of the antibodies. The primary antibodies used were against α-smooth muscle actin (Abcam, ab7817, 1:300), mannose receptor (Abcam, ab64693, 1:100), myocardial related transcription factor A (Santa Cruz, sc-398675, 1:100), and Rho associated coiled-coil containing protein kinase 2 (Santa Cruz, sc-398519, 1:100). These antibodies were labeled with fluorophore-conjugated, anti-IgG antibodies (Thermo Fisher, AlexaFluor, 1:400) to enable their detection. The nuclei were counterstained with Hoechst 33342 (Thermo Fisher, 1:1000).

To measure the fluorescence intensity of α-SMA in different microtissues under varied pharmacological conditions, images were taken on a Nikon Ti-U microscope equipped with a Hamamatsu ORCA-ER CCD camera using a 10× Plan Fluor objective under identical imaging conditions. Fluorescence intensity was measured in ImageJ by taking the integrated intensity of a region of interest bounded by the inner edge of micropillars and then subtracting the background intensity. Confocal images of the microtissues were taken on a Leica Stellaris 5 fluorescence microscope with objectives (10x air objective and 40x oil immersion). An optical slice of 1 μm was used for all channels, and the stack of images was processed using the Z stack tool in Leica LAS X software package.

The number of macrophages remaining attached on the surface of microtissues, and glass coverslips was quantified from single color fluorescence images using ImageJ, a software program developed by the U.S. National Institutes of Health (NIH), Bethesda, MA, USA, and available for free download [http://imagej.nih.gov/ij/; 1997–2013].

### Histologic analysis of lung sections

Human lung tissue samples from patients diagnosed with IPF were obtained from the NIH tissue bank. The tissue was cut into small pieces measuring 0.3 cm to 0.5 cm in one dimension and embedded in Tissue-Tek® O.C.T. Compound (4583, Sakura Finetek). The embedded tissue was rapidly frozen on dry ice and cut into 15 μm cryostat sections. The sections were then mounted on microscope slides (MS-CS-90, 45o Corners, stellar scientific) and subjected to immunofluorescent staining and H&E staining.

### Scanning electron microscopy

The microdevices containing normal human lung fibroblasts with or without macrophages were fixed using a solution of 2% glutaraldehyde (ACROS organics) in PBS (containing Ca2+/Mg2+) for 1 h at 4°C. Next, the samples were dehydrated with ethanol in increments of 10% from 50% to 100% for 10 min each, followed by critical point drying for 1 hour using hexamethyldisilazane (HMDS, Alfa Aesar) in a chemical hood. The scanning electron microscopy (SEM) was performed using the Hitachi SU70 Field Emission Scanning Electron Microscope (FESEM).

### Single cell RNA-seq data mining

To obtain ITGAM, ITGB2, FN1, and RHOA gene expression data for human lung tissue, NCBI GEO database GSE122960 and GSE136831 were accessed. In the analysis of GSE122960, a two-tailed unpaired t-test was conducted using GraphPad Prism version 8.0 for Windows (GraphPad Software, www.graphpad.com). The scRNA-seq data from GSE122960 was visualized using UCSC CellBrowser, developed by Maximilian Haeussler (https://www.nupulmonary.org/resources/). Additionally, the scRNA-seq data from GSE136831 was visualized using Cell Explorer in the IPF Cell Atlas (http://www.ipfcellatlas.com/).

To generate the t-distributed stochastic neighbor embedding (t-SNE) plots for alveolar macrophages (Fig.1 C, D), cells identified as macrophages through individual annotation of single-cell RNA-Seq data from eight normal lungs and eight fibrotic lungs were aggregated. On the tSNE plot representing all 16 samples, cells were categorized based on their origin as either coming from a donor or from a patient with pulmonary fibrosis. The color scheme was used to differentiate between these two groups.

### Statistical analysis

All experiments were conducted with a minimum of three biological replicates. The quantitative data are presented as means ± standard deviation (SD). Statistical differences between groups were evaluated using analysis of variance (ANOVA) followed by a post hoc Tukey’s multiple comparisons test and Dunnett’s multiple comparisons test. The significance levels were set at *P = 0.05 and **P = 0.01. For experiments comparing two groups, a two-tailed paired Student’s t-test was performed. Statistically significant differences were denoted as *P ≤ 0.05, **P ≤ 0.01, and ***P ≤ 0.001.

## Acknowledgments

We would like to express our gratitude to Dr. Daniel B. Rifkin from New York University Grossman School of Medicine for generously providing TMLCs. We also thank Lisa Eagler at the Clinical and Translational Research Center at University at Buffalo for her assistance with blood draw from healthy volunteers. We acknowledge Dr. Pedro Lei at the Cell, Gene, and Tissue Engineering Center at University at Buffalo for providing technical support on the Leica Stellaris 5.0, and Dr. Peter Bush at South Campus Instrument Center at University at Buffalo for capturing SEM images.

## Author contributions

Y.X. and R.Z. conceived the idea and designed the experiments; Y.X., LX.Y., J.K.L. performed experiments; Y.X. analyzed data; Y.X. and R.Z. wrote the manuscript; B.H., J.K.L., Y.X. and R.Z. edited the manuscript; and R.Z. provided financial and administrative support.

## Data and materials availability

All data needed to evaluate the conclusions in the paper are present in the paper and/or the Supplementary Materials. Additional data related to this paper may be requested from the authors.

Uncategorized References

