## supplementary figures for "Modeling Mechanical Activation of Macrophages During Pulmonary Fibrogenesis for Targeted Anti-Fibrosis Therapy"

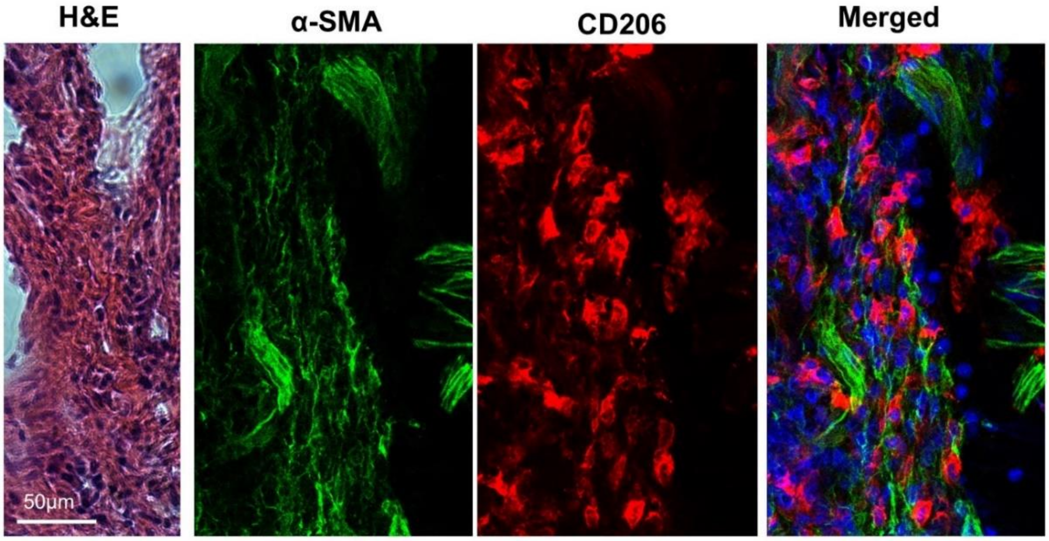


**SFig. 1** (A) H&E staining of a portion of fibrotic lung tissue. (B) Confocal fluorescence images of an adjacent tissue slice of the samples shown in (A). Myofibroblast marker α-SMA (green), macrophage marker CD206 (red) and the merged channels with cell nuclei stained with 4′,6-diamidino-2-phenylindole (DAPI)(blue). Scale bars, 50 μm.

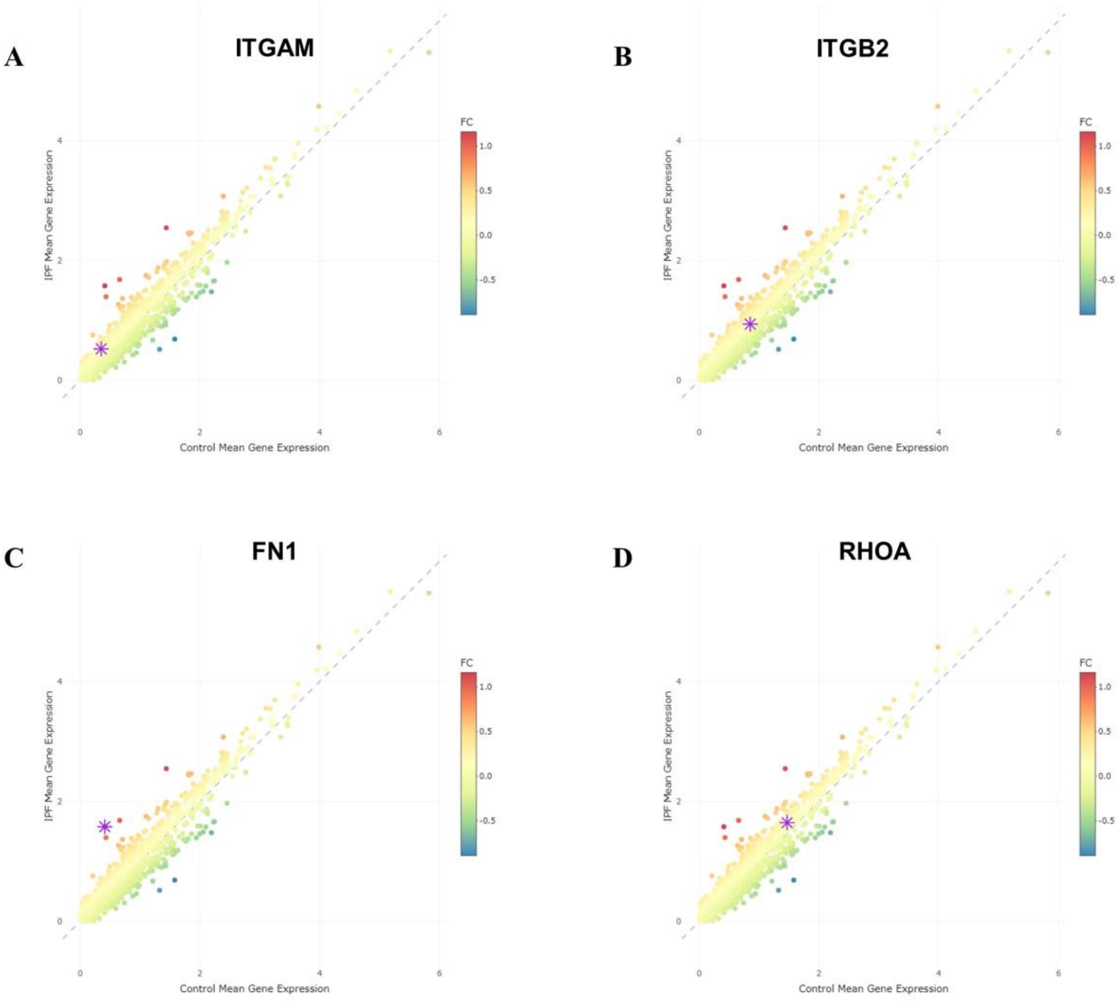


**SFig. 2** A scatter plot of the average of gene expression of **(A)** ITGAM, **(B)** ITGB2, **(C)** FN1 and **(D)** RHOA in healthy (x-axis) and IPF (y-axis) lung samples. Points are colored by their fold ratios; progressive shades of red indicate increase, and progressive shades of blue indicate decrease. Gray dashed lines are lines that depict zero-fold change. Data extracted from 312,928 cells from distal lung parenchyma samples obtained from 32 IPF, 18 COPD and 29 healthy (control) donor lungs based on NCBI GEO Dataset: GSE136831.

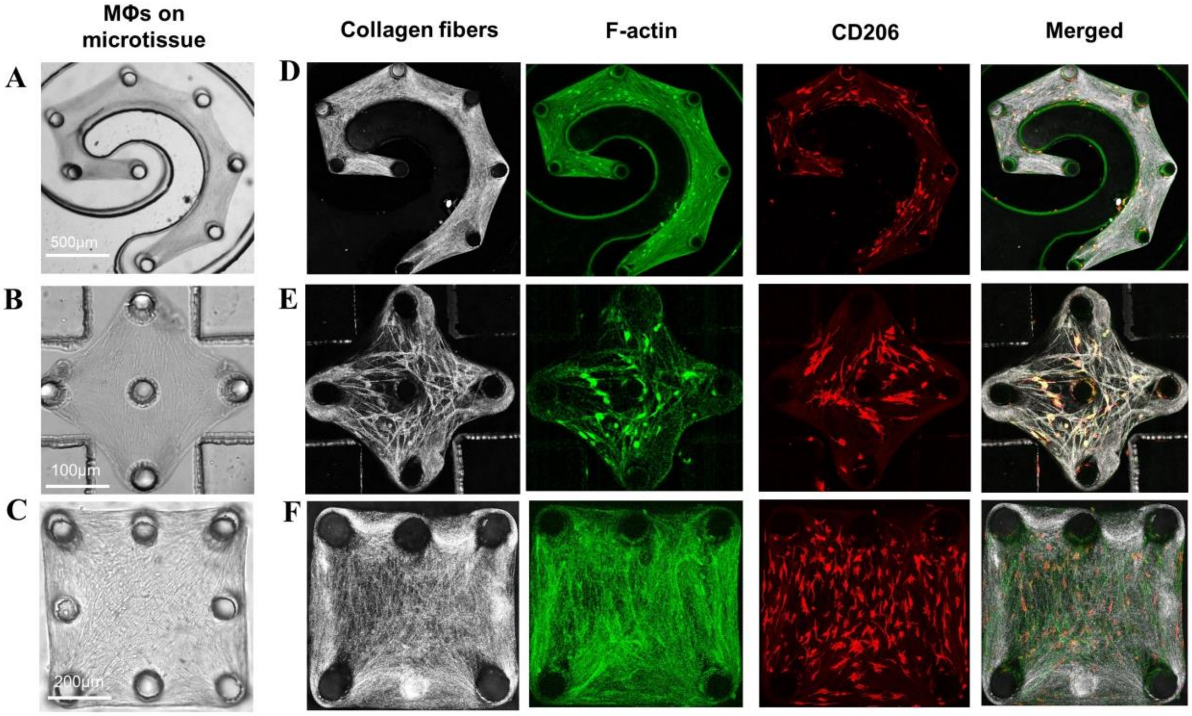


**SFig. 3** (A, B, C) Phase contrast images of fibroblast and macrophage co-cultured microtissues. (D, E, F) Confocal reflectance images of collagen fibers (gray), and fluorescence images of F-actin (green), CD206 (red) and merged channels.

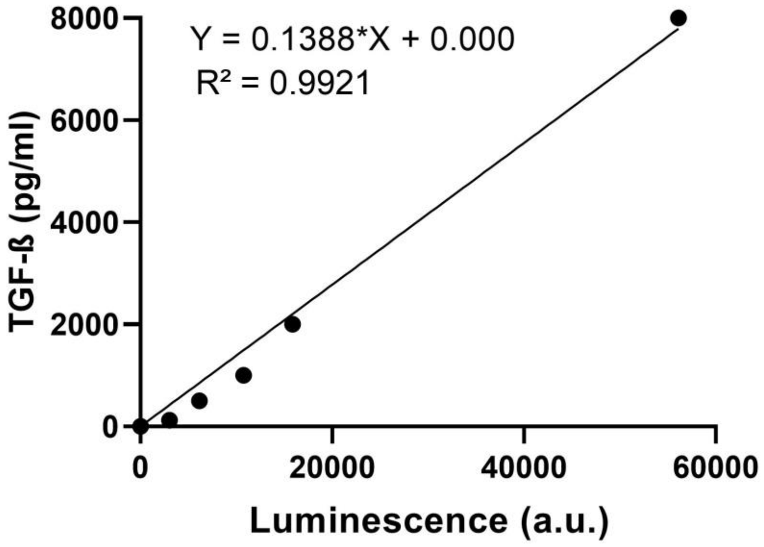


**SFig. 4** A representative standard curve for active TGF-β measurement using TMLC reporter cells. Luminescence measurements were converted into ng/ml of TGF-β by establishing standard curves performed with known concentrations of active TGF-β.

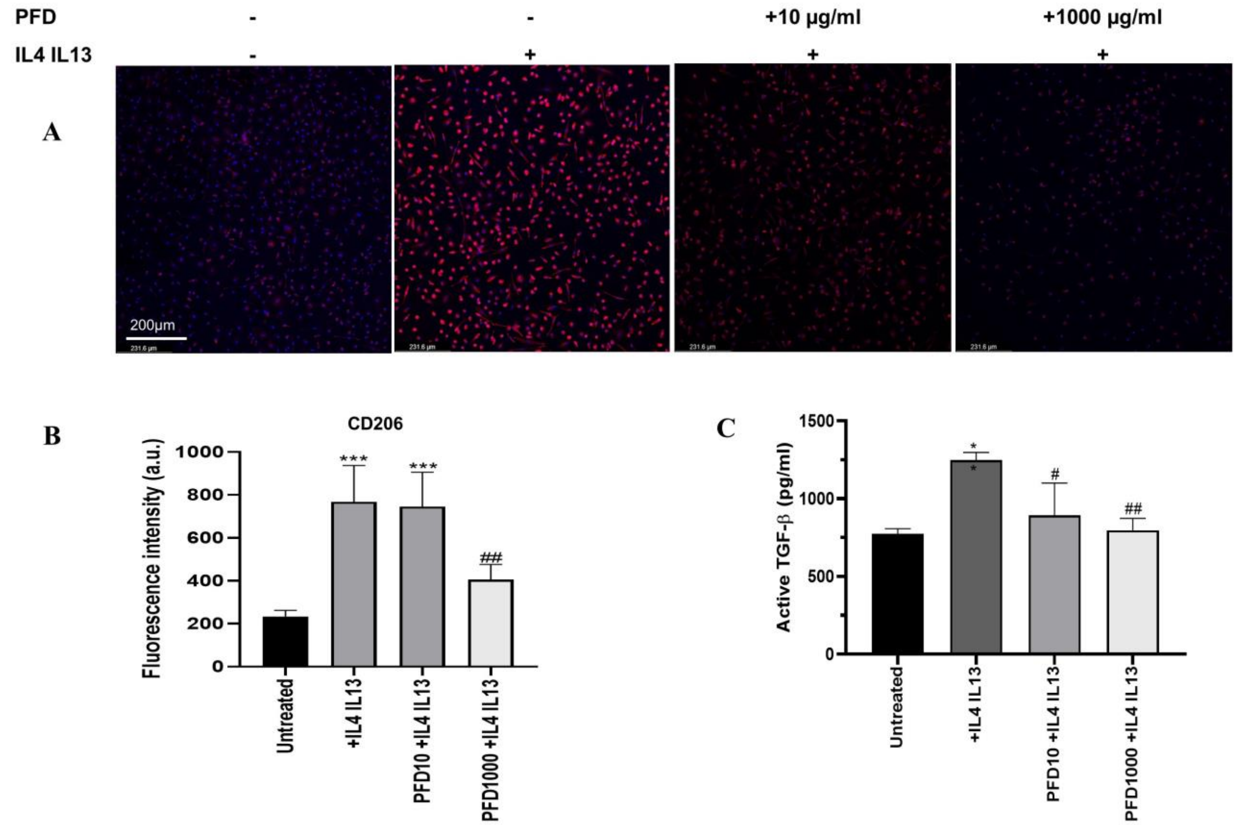


**SFig. 5** Effects of Pirfenidone (PFD) on the polarization of human PBMC derived monocytes towards the M2 phenotype. (A) Immunofluorescence staining of the M2 phenotypic marker CD206 (red) in monocytes pretreated with PFD for 24hs followed by the stimulation with or without IL-4 (20 ng/ml) + IL-13 (20 ng/ml) for additional 24hs. (B) Fluorescent intensity of CD206 in macrophages treated without or with PFD. (C) The concentrations of active TGF-β in cell culture supernatant of different culture conditions. Data are means ± SD from at least five experiments. *p<0.05 and ***p<0.01, when compared with the untreated group. #p<0.05 and ##p<0.01, when compared with the IL-4 + IL-13 stimulated group.

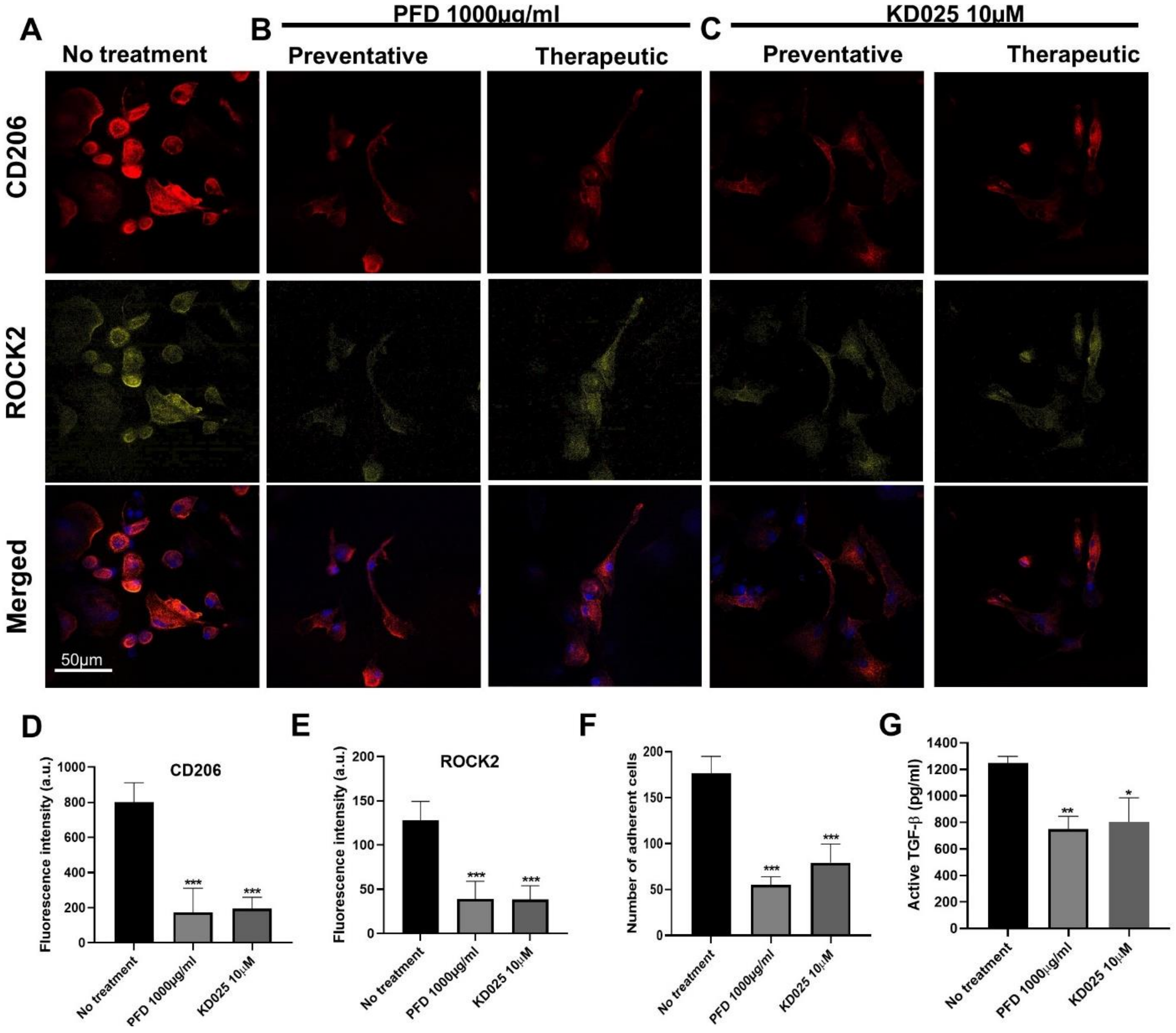


**SFig. 6** Representative immunofluorescence images of monocultured macrophages under no treatment (A), PFD (B) or KD025 (C) treatments. Cells were seeded on PDMS with stiffness of 4 MPa and stained for CD206 (red), ROCK2 (yellow) and Nuclei (blue) from top to bottom. Fluorescent intensity of CD206 (D) and ROCK2 (E) in monoculture macrophages treated without or with PFD or KD025. (F) Number of adherent macrophages after treatments. (G) The concentrations of active TGF-β in cell culture supernatant of different treatment conditions.

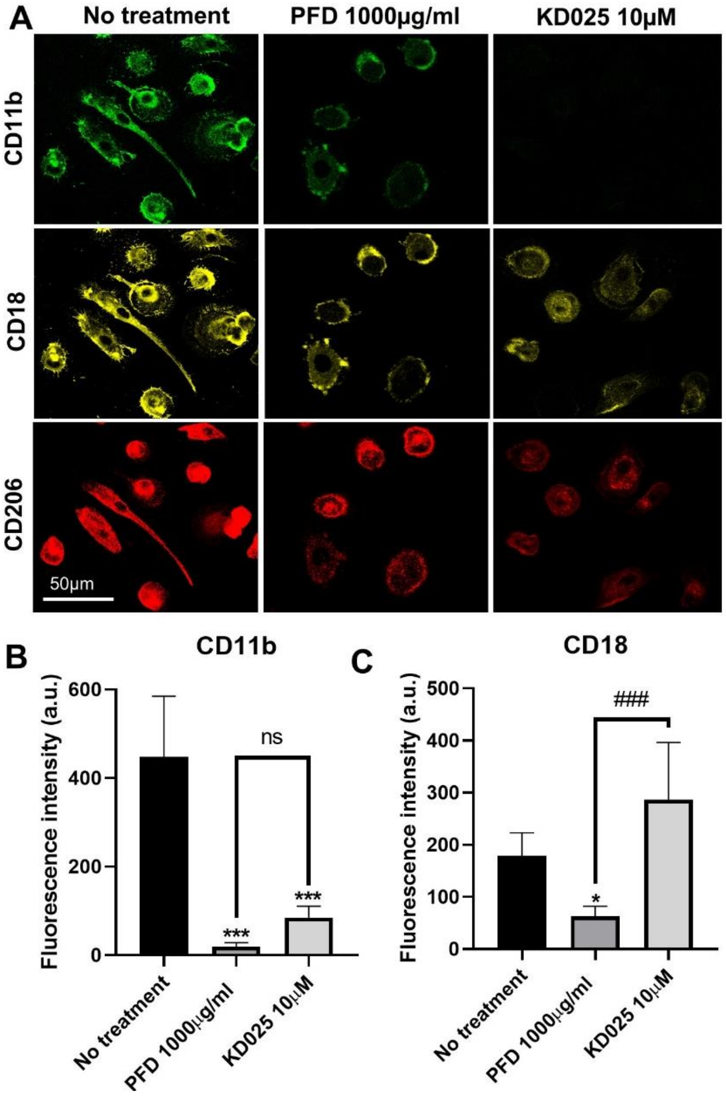


**SFig. 7** (A) Representative immunofluorescence images of monocultured macrophages under PFD or ROCK2 inhibitor KD025 treatments. Cells were seeded on glass and stained for CD11b (green), CD18 (yellow) and CD206 (red) from top to bottom. Fluorescent intensity of CD11b (B) and CD18 (C) in macrophages treated without or with PFD or KD025.
